# EEG-based Schizophrenia Detection Using Spectral, Entropy, and Graph Connectivity Features with Machine Learning

**DOI:** 10.64898/2026.04.08.717137

**Authors:** Nazila Ahmadi Daryakenari, Seyed Kamaledin Setarehdan

## Abstract

Schizophrenia is a serious mental disorder that changes the way people think, perceive, and manage daily life. Getting the diagnosis right is critical for proper treatment, but in practice it is often difficult. Current evaluations depend mostly on a clinician’s judgment, and the overlap of symptoms with bipolar disorder or major depression makes the task even harder. EEG offers a safe and noninvasive way to study brain activity, yet no single EEG feature has been reliable enough to stand on its own. This makes it important to look at integrative approaches that bring together different aspects of brain dynamics.

In this study, we analyzed EEG features to distinguish patients with schizophrenia from healthy controls. Spectral power was measured across δ, θ, α, β, and γ bands. Temporal irregularity was quantified with Multiscale Permutation Entropy (MPE), which to our knowledge represents the first application of MPE to EEG in schizophrenia. Functional connectivity was estimated with the weighted Phase Lag Index in θ, α, and β bands, followed by extraction of graph measures including global efficiency, clustering coefficient, characteristic path length, and mean strength. These features were used to train Random Forest, Multi-Layer Perceptron, and Support Vector Machine classifiers. Among the models, Random Forest achieved the most reliable performance, reaching 99.7% accuracy under stratified 5-fold validation and 99.6% under leave-one-subject-out validation. Feature analysis showed that connectivity in θ and α bands contributed most strongly to classification. Topographic maps of θ, α, and β activity also revealed regional group differences. Overall, the results suggest that combining spectral, entropy, and connectivity measures offers a promising framework for EEG-based detection of schizophrenia. Nevertheless, these findings are preliminary given the limited sample size (N=28), and replication in larger and more diverse cohorts is required before clinical translation.

## I. Introduction

Schizophrenia is a long-term psychiatric illness that profoundly alters the way people think, feel, and behave. It often leads to social withdrawal, a loss of motivation, and a flattening of emotional expression. About one percent of the world’s population is affected, making it a major public health challenge. At present, diagnosis still relies mostly on clinical interviews and standardized checklists such as DSM-5 or ICD-10 [1]. In the absence of reliable biological markers, patients are frequently misdiagnosed or diagnosed late, and research cohorts often include highly heterogeneous cases. This situation makes the search for measurable, brain-based markers an urgent clinical priority.

The global burden of schizophrenia is considerable. According to the World Health Organization (WHO), about 26 million people live with the disorder worldwide, and the annual cost of treatment is estimated at 32.5–65 billion USD. The human cost is also devastating: around 5% of patients die by suicide. Symptoms most often emerge in late adolescence or early adulthood, developing gradually over time. The condition tends to appear earlier in men than in women and is more common in urban populations and minority groups [2]. Even so, WHO stresses that schizophrenia is treatable, and diagnosis—whether early or late—remains critical for guiding appropriate care and improving outcomes.

Access to accurate diagnosis, however, is highly uneven. Nearly 90% of untreated patients live in low- and middle-income countries, where mental health resources are limited. Traditional diagnosis requires long and costly interviews carried out by trained psychiatrists, a process that is not only exhausting but also prone to human error. This underscores the need for automated systems that are accurate, affordable, and capable of supporting clinicians in making faster and more reliable decisions [3].

Electroencephalography (EEG) offers a promising solution. It is inexpensive, widely available, and captures brain activity with millisecond precision. Compared to other neuroimaging techniques such as functional MRI, EEG is more flexible in design, less costly, and offers far superior temporal resolution. These advantages have led to growing interest in EEG for the study and classification of schizophrenia. As a non-invasive and practical tool, EEG-based diagnosis has the potential to improve early and accurate detection of the disorder and to serve as a valuable complement in clinical practice [4].

Extensive research has examined EEG-based features to distinguish individuals with schizophrenia (SZ) from healthy controls (HC). Resting-state EEG investigations have reported shifts in power across frequency bands [5]. Patients with SZ often show increased power in lower ranges such as delta and theta, and reduced power in higher bands including alpha, beta, and gamma [6, 7].

Multiscale characteristics are a common property of time series generated by complex systems [8]. The dynamic behavior of such systems can be more effectively captured when multiple temporal scales are considered [9]. Multiscale Entropy (MSE), as proposed by Costa et al. [10, 11], has been shown to reveal structural information embedded in different scale factors of time series [11, 12]. Building on this idea, Bai et al. [13] identified schizophrenia using multiscale recurrence information in MEG signals. In a recent study, Dengxuan Bai et al. [14] utilized Multiscale Weighted Permutation Entropy, a method based on entropy theory, to detect schizophrenia through MEG data. These findings underscore the potential of analyzing multiscale properties of time series to offer more profound insights into the complexity underlying brain dynamics in schizophrenia.

Using indices like coherence and the Phase Lag Index (PLI), researchers have shown widespread dysconnectivity in SZ, particularly in the theta, alpha, and beta ranges [15, 16]. EEG-derived networks constructed with weighted PLI often reveal reduced global efficiency and clustering, pointing to impaired integration and segregation of brain activity. Despite these insights, a common limitation in the literature is that spectral, complexity, and connectivity measures are usually studied separately.

To address these shortcomings, we present a multi-domain EEG analysis framework for schizophrenia classification. The novelty of our work lies in three elements. First, we integrate spectral, complexity, and connectivity features within a unified EEG framework. We simultaneously extract features from all three domains: band power from delta to gamma, and functional connectivity via Weighted Phase Lag Index. From the resulting networks, we compute graph metrics including global efficiency, clustering coefficient, characteristic path length, and mean strength. Specifically, we implement Multiscale Permutation Entropy (MPE) across six temporal scales. It has previously been explored in magnetoencephalography (MEG) and other neuroimaging modalities, but, to our knowledge, the present study is the first to apply MPE to EEG data in the context of schizophrenia classification. This extension is important because EEG is more accessible and widely used than MEG, suggesting that entropy-based metrics may hold promise as practical biomarkers. Secondly, a rigorous evaluation of multiple machine learning classifiers such as Random Forest, Support Vector Machine, and Multi-Layer Perceptron classifiers is conducted under two validation schemes: stratified 5-fold cross validation and leave-one-subject-out cross validation. The latter provides a clinically realistic estimate of generalizability. Thirdly, we advance beyond mere accuracy by conducting a comprehensive analysis of feature importance within the Random Forest model. This approach enables the identification of the most discriminative neural signatures, including connectivity patterns in the theta band. Beyond classification accuracy, we use feature importance analysis within the Random Forest framework to highlight which neural signatures contribute most to group separation. This provides interpretable insights into the neural basis of SZ, such as the role of theta-band connectivity, and informs future biomarker development.

We expect this integrative strategy to deliver more reliable classification performance than approaches confined to a single feature type. The findings aim not only to improve discrimination between patients with schizophrenia and healthy controls but also to support the development of interpretable and clinically meaningful EEG-based biomarkers.

## II. Dataset and Methods

### A. Dataset

The EEG dataset was recorded at the Atieh Clinical Neuroscience Center. It included 14 healthy participants and 14 patients diagnosed with schizophrenia, with mean ages of 25.3 ± 8.1 and 28.4 ± 5.9 years, respectively. Signals were sampled at 500 Hz using 19 electrodes placed according to the international 10–20 system. Recordings were made during rest with eyes closed for 5–7 minutes.

### B. Preprocessing

Electroencephalographic (EEG) recordings from patients diagnosed with schizophrenia (SZ) and from healthy controls (HC) were subjected to rigorous preprocessing in MNE. Signals were filtered with a fourth-order Butterworth band-pass (1–45 Hz) and re-referenced to the average. The continuous recordings were segmented into two-second windows with 50% overlap: this length balances temporal stationarity and spectral resolution, while the overlap increases the number of epochs and prevents transient patterns from being missed at window boundaries. Artifact handling combined automatic and ICA-based approaches: independent components correlated with ocular or cardiac activity were excluded, muscle-related bursts were annotated automatically, and epochs exceeding conservative amplitude thresholds (∼120–150 µV) were rejected. Visual inspection was used only for quality control, with all rejection thresholds fixed a priori to avoid bias.

### C. Feature Extraction

#### 1) Spectral power

A conventional approach to analyzing electroencephalogram involves the decomposition of the signal into distinct frequency bands, namely delta (1–4 Hz), theta (4–7 Hz), alpha (8–12 Hz), beta (13–30 Hz), and gamma (30–45 Hz). Schizophrenia has been reported to affect activity across all of these bands. To quantify band-specific activity, we first estimated the power spectral density (PSD) using Welch’s method, which averages Fourier transforms of overlapping signal segments to produce a stable spectrum. The mean band power was then obtained by integrating PSD values within each frequency band. Finally, channel-wise values were averaged to produce one measure per band for each epoch [17].

#### 2) Multiscale permutation entropy

To quantify temporal irregularity, we employed Multiscale Permutation Entropy (MPE). Unlike single-scale entropy measures, MPE evaluates complexity across multiple temporal resolutions. To compute MPE, three main steps are required: (i) coarse-graining of the original time series at different scales, (ii) calculation of permutation entropy for each scale, and (iii) construction of the multiscale feature vector.

##### a) Coarse-graining

Given a one-dimensional time series {*x*(*i*) : *i* = 1,2, . ., *N*}, the multiscale process begins by constructing a coarse-grained series. Consecutive samples are averaged in non-overlapping groups of length s, generating a coarse-grained sequence *y*_*j*_^(*s*)^ that reflects slower temporal dynamics of the original signal. When the scale factor is s=1, the coarse-grained sequence is identical to the original series [18]. For a general scale s, the coarse-grained time series is defined as:

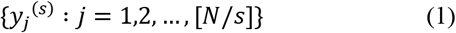

where each element is calculated as:

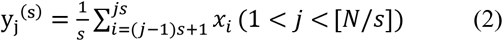

##### b) Permutation entropy at scale s

For each coarse-grained series, permutation entropy is calculated. Using embedding dimension *m*, ordinal patterns are constructed and their probabilities *p*_*i*_^(*s*)^ are obtained. The normalized permutation entropy is defined as:

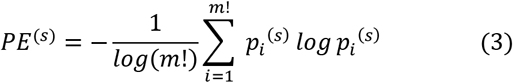

This quantifies the irregularity of the signal at the chosen scale[11].

##### c) Multiscale feature vector

Repeating the above steps for scales S =1,2, …,s yields the final multiscale feature vector:

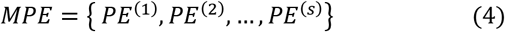

In this study, six scales were used (S=6), enabling the characterization of both fast and slow temporal dynamics.

#### 3) Functional connectivity and graph measures

The estimation of functional connectivity between EEG channels was performed using the weighted Phase Lag Index (wPLI), a method that quantifies consistent phase relationships while reducing the effect of volume conduction [19]. WPLI utilizes a cross-spectrum evaluation method that prioritizes the magnitude of the imaginary component. This allows it to limit the influence of cross-spectrum elements around the real axes which are at risk of changing their “true” sign with small noise perturbations. For two signals *x*_*i*_(*t*) and *x*_*j*_(*t*), wPLI is defined as:

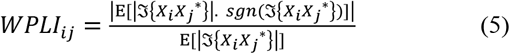

where *X*_*i*_ is the analytic signal of channel i obtained via the Hilbert transform, and 𝔍 denotes the imaginary part. From the resulting connectivity matrices, graph-theoretical metrics were computed to characterize network topology [20, 21]:

##### a) Global efficiency

Global efficiency quantifies how efficiently information is exchanged across the entire network. It is defined as the average inverse shortest path length:

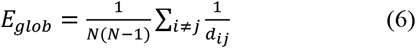

where *d*_*ij*_ is the shortest path length between nodes i and j. Lower efficiency in schizophrenia suggests impaired integration of distributed brain regions.

##### b) Clustering coefficient

The clustering coefficient is indicative of the tendency of nodes to form tightly interconnected groups, thereby indicating local specialization. For weighted networks, the given formula is as follows:

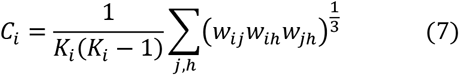

Where *K*_*i*_ is the degree of node i and *w*_*ij*_ is the weight of edge (i,j). Reduced clustering in individuals diagnosed with schizophrenia has been demonstrated to indicate a weakening of segregation within local subnetworks.

##### c) Characteristic path length

This metric represents the mean shortest path length across all node pairs:

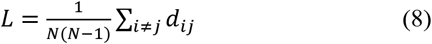

Higher values imply reduced communication efficiency across the network. Increased path length has been linked to less efficient global information transfer in individuals diagnosed with schizophrenia.

##### d) Mean strength

Node strength is the sum of weights of edges connected to a node, and mean strength is the average across all nodes:

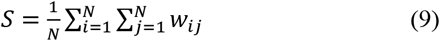

It represents the overall level of functional connectivity. The observed decrease in mean strength among patients indicates a potential for weaker global coupling among brain regions.

### D. Classification of Features

Compared three widely used machine learning models: Random Forest (RF), Support Vector Machine (SVM), and Multi-Layer Perceptron (MLP). The Random Forest was built with 500 trees using the Gini split criterion. We allowed the trees to grow fully (unrestricted depth), but quick tests with limited depth (10–15 levels) gave almost identical results, showing that the accuracy was not just a by-product of tree growth. The MLP had two hidden layers with 64 and 32 neurons, used ReLU activations, and a sigmoid output layer. It was trained with the Adam optimizer (learning rate 0.001, up to 500 iterations). Before entering the models, all features were standardized. Hyperparameter settings followed prior EEG studies and were further confirmed with compact grid search. Analyses were performed in Python 3.13.5 using scikit-learn 1.7.1, MNE-Python 1.10.1, NumPy 2.3.1, and SciPy 1.16.0.

#### 1) Random Forest (RF)

Random Forest is an ensemble learning method that builds many decision trees and combines their predictions. Each tree is trained on a random sample of the data, and at every split, a random subset of features is considered. This randomness creates diversity across trees and helps the model avoid overfitting, leading to strong generalization. Random Forest is well suited for high-dimensional EEG data because it can capture complex nonlinear patterns without heavy parameter tuning. An additional strength is its ability to rank feature importance, providing interpretable insights into which features drive classification [22]. Compared with the other models, RF often provides the most stable and generalizable results, even when data are noisy or high-dimensional.

#### 2) Linear Support Vector Machine

Support Vector Machines are widely used for supervised classification. Their goal is to find the best hyperplane that separates two classes in a high-dimensional space. The model works by maximizing the margin between classes, which improves generalization. Only a small number of training points, called support vectors, determine the position of this hyperplane. These critical points sit closest to the boundary and define the decision rule [23]. Compared with RF and MLP, SVMs usually perform best when the dataset is not very large and when the boundary between classes is sharp.

#### 3) Multi-Layer Perceptron

The Multi-Layer Perceptron is a type of feedforward neural network used for tasks such as classification [24]. It consists of an input layer, one or more hidden layers, and an output layer. Each layer is made up of neurons that transform the input they receive and pass it to the next layer. The network is fully connected, meaning every neuron in one layer connects to every neuron in the next. Information flows through these connections with associated weights, which determine the strength and influence of the signals being transmitted [25]. Compared with RF and SVM, the MLP can model more complex relationships, but it may require more data and careful tuning to reach its full potential.

### E. Validation

#### 1) Stratified K-fold cross-validation

The evaluation of classifier performance requires the implementation of reliable validation strategies. One of the most common approaches is k-fold cross-validation, in which the dataset is divided into k partitions. The dataset is partitioned into K folds, with each fold utilized once for the testing stage. The remaining folds are then employed for the training phase. A potential limitation of simple k-fold partitioning is that it may not preserve the balance between classes across folds, which can result in biased evaluations. To address this, we employed stratified k-fold cross-validation, where each fold is created so that the proportion of schizophrenia patients and healthy controls matches the original distribution. This approach ensures that both classes are represented consistently in all folds, thereby providing a more robust estimate of classifier performance [26].

In the present implementation, stratified 5-fold cross-validation was applied at the subject level. This approach ensures that all epochs from the same individual are maintained within the same fold, thereby preventing data leakage between the training and test sets. Despite the balanced sample size of 14 patients and healthy controls in our dataset, we implemented stratified k-fold cross-validation to ensure the preservation of class proportions in each fold. This approach aligns with established best practices in validation design.

#### 2) Leave-one-subject-out

In addition to stratified 5-fold cross-validation, we employed leave-one-subject-out (LOSO) validation. With this method, data from one subject is withheld for testing while the model is trained using all the remaining subjects. This process is repeated until each subject has been used as the test set once. LOSO provides a stricter and more clinically meaningful evaluation since the model is tested on entirely unseen individuals rather than segments of subjects already in the training set. This design reduces the risk of data leakage at the subject level and provides a more realistic estimate of generalization performance in real-world applications [27].

To make the workflow clearer, we added a methodological flowchart (Fig. 1). It shows the pipeline step by step, from EEG acquisition and preprocessing to feature extraction and classification. The diagram also illustrates how spectral, entropy, and graph connectivity features were brought together before being used in the machine learning models.

**Fig. 1.**
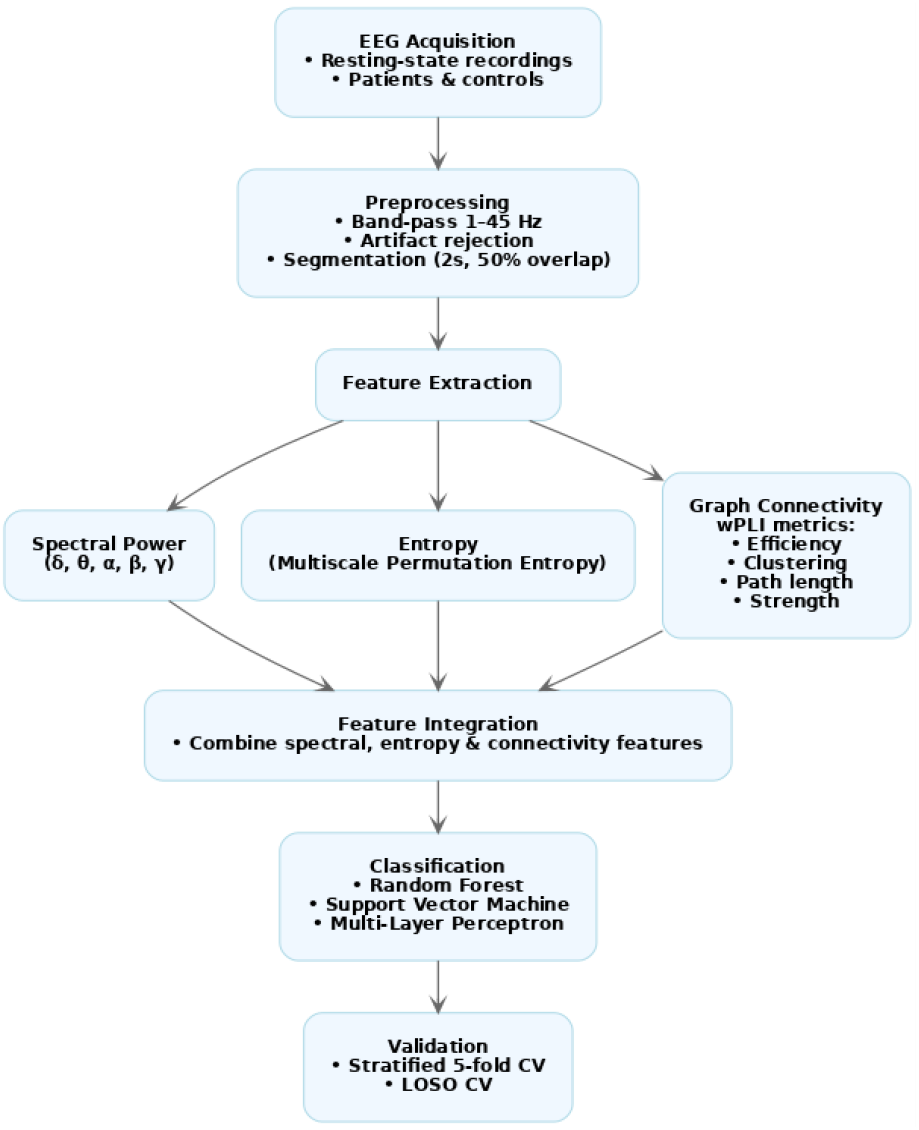
Methodological flowchart of the proposed EEG-based schizophrenia detection pipeline.

## III. Results and Discussion

### A. Multiscale permutation entropy

Multiscale permutation entropy (MPE) analysis revealed significant differences between patients with schizophrenia and healthy control subjects. On average, patients exhibited higher entropy values across six temporal scales, as well as greater variability from scale to scale. The mean MPE was 0.49 ± 0.01 for patients and 0.43 ± 0.02 for controls, and the respective standard deviations were 0.44 ± 0.02 and 0.39 ± 0.02. These results suggest that EEG dynamics are more irregular and less predictable in schizophrenia, indicating an unstable temporal organization of brain activity.

The spatial distribution of these differences added further insight. Frontal electrodes (Fp1, F7, Fz, F8) showed the strongest increases in mean entropy, with additional changes along central–parietal midline sites and in occipital regions. Variability across scales (MPE-std) was also most pronounced at frontal (F3, F4), temporal (T7, T8), and parietal (P3, P4) electrodes, pointing to abnormal fluctuations that are regionally distributed rather than global.

Taken together, these results suggest that schizophrenia is marked by abnormal neural complexity across cognitive and sensory systems. Increased entropy in frontal regions points to impaired executive control and network disorganization, while occipital and temporal changes align with instability in visual and auditory pathways. This pattern highlights schizophrenia as a disorder of disrupted temporal complexity that extends beyond a single cortical region. By capturing these multiscale alterations, MPE may offer a clinically accessible biomarker to support diagnosis and guide future research on the neural basis of schizophrenia.

These EEG results extend previous MEG studies, where both multiscale recurrence entropy and multiscale weighted permutation entropy distinguished schizophrenia patients from healthy participants [13, 14]. To our knowledge, this is among the first demonstrations of MPE applied to EEG in schizophrenia, offering a clinically accessible way to capture complexity changes. Importantly, the multiscale approach reveals disturbances that are not always visible in single-scale entropy analyses, which have sometimes reported inconsistent findings [21].

Fig. 2 illustrates the spatial distribution of these differences, showing frontal peaks in mean MPE and frontal–temporal peaks in variability, thereby providing a spatial view of how multiscale complexity is altered in schizophrenia.

**Fig. 2.**
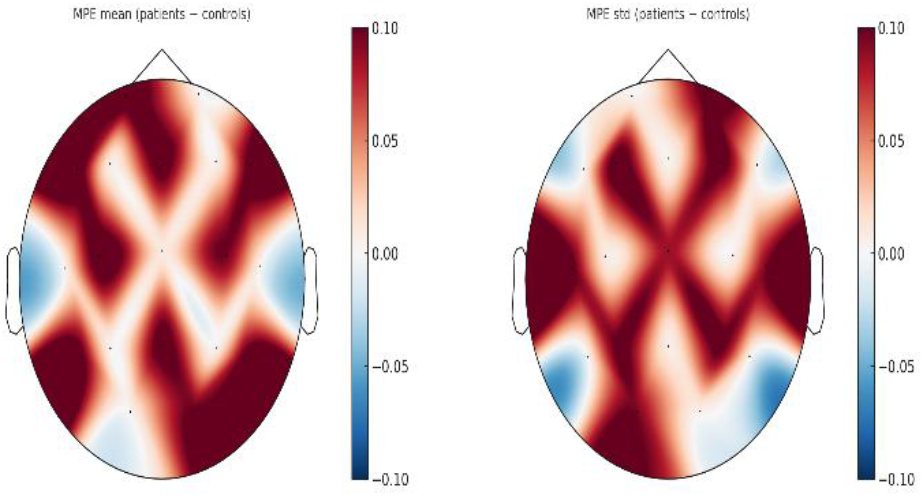
Topographical maps of multiscale permutation entropy (MPE) differences. Left: Mean MPE (patients − controls). Right: Standard deviation across scales, with the largest increases over frontal–temporal regions.

### B. Spectral Power Topography

Topographical analysis of EEG band power revealed consistent abnormal activity patterns in patients with schizophrenia compared to healthy controls. we found a pronounced increase in low-frequency activity (Delta and Theta bands) alongside a marked decrease in high-frequency activity (Alpha, Beta, and Gamma bands).

Delta power typically increases most prominently over frontal-central regions, reflecting cortical hypoarousal and thalamo-cortical dysregulation. Elevated theta activity is most evident at midline frontal sites (ACC and mPFC) and is associated with deficits in cognitive control and working memory [28]. Alpha power is consistently reduced, especially in frontal–parietal regions, indicating weaker network synchronization and impaired inhibitory control [29]. Resting-state beta findings are mixed. Some studies report frontal–central reductions, but the effects are often weak. Task-based studies show more consistent abnormalities, such as reduced beta desynchronization during emotional and motor processing [29, 30]. Gamma power is broadly reduced, particularly in temporal-parietal regions, and is linked to GABAergic interneuron dysfunction. However, recent meta-analyses show heterogeneous results (increases, decreases, or null findings). Elevated frontal gamma power has also been associated with greater positive symptom severity [31, 32].

Table I summarizes the topographic results, showing a clear pattern of stronger low-frequency activity and weaker high-frequency activity across brain regions in schizophrenia. Fig. 3 illustrates these differences on topographical maps, giving a spatial view of where the changes between patients and controls are most pronounced.

**TABLE I.** Summary of EEG band power alterations in schizophrenia.

| Frequency Band | Observed in our study | Most affected regions | Reported in literature | Associated mechanisms |
| --- | --- | --- | --- | --- |
| Delta ($\delta$) | Increased power | Frontal and central | Elevated, especially fronto-central | Cortical hypoarousal; thalamo-cortical dysregulation |
| Theta ($\theta$) | Increased power | Fronto-midline | Elevated, often at midline | Altered thalamo-cortical loops; deficits in cognitive control |
| Alpha ($\alpha$) | Decreased power | Frontal and centro-parietal | Reduced across frontal–parietal, sometimes occipital | Weakened large-scale synchronization; reduced inhibitory regulation |
| Beta ($\beta$) | Weak reduction (not robust) | Diffuse, no strong focal effects | Decreased over frontal–central (variable across studies) | Altered motor and attentional networks |
| Gamma ($\gamma$) | Broad reduction | Temporal and parietal | Consistently reduced, especially temporal–parietal | GABAergic interneuron dysfunction; impaired perceptual binding |

**Fig. 3.**
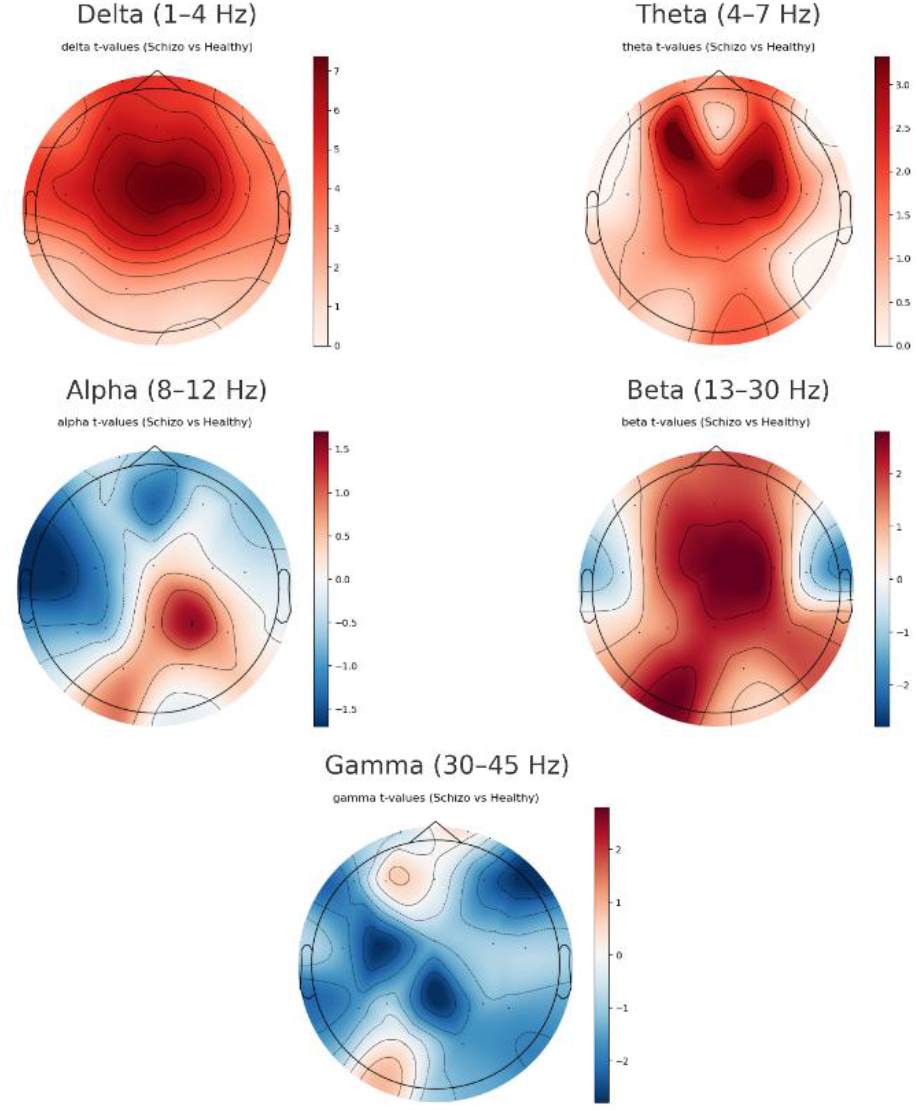
Topographical maps of EEG band power differences between patients with schizophrenia and healthy controls. Delta and theta are increased fronto-centrally, while alpha and gamma show widespread reductions.

### C. Connectivity and Graph Measures

When we looked at the brain networks using graph measures, the patients with schizophrenia showed a clear loss of efficiency. The global efficiency of their networks was lower, which means that information could not travel as smoothly between distant brain regions. The clustering coefficient was also reduced, pointing to weaker local groupings of neurons that normally work together. At the same time, the path length was longer, so signals needed more steps to move from one region to another. Finally, the overall strength of connections was reduced, showing that synchronization between brain areas was simply weaker.

Taken together, this suggests that the brain networks of patients have shifted toward a more random and less optimized structure, moving away from the balanced “small-world” pattern that healthy brains usually have. This reorganization offers a good explanation for many of the cognitive difficulties seen in schizophrenia, such as slower thinking, reduced memory capacity, and disorganized thought.

The wPLI maps give a spatial picture of these changes. Connectivity was broadly reduced, but the drop was especially strong in the alpha and beta bands, rhythms that are important for attention, working memory, and higher-order integration. Frontal and temporal regions were most affected, which fits well with the idea of hypofrontality and the long-standing view that schizophrenia disrupts the brain’s key control hubs. Theta-band reductions were also present, particularly over midline areas, pointing to problems with cognitive control.

These results are very much in line with what recent studies and meta-analyses have reported. Research up to 2025 consistently shows the same pattern of reduced efficiency, lower clustering, longer path lengths, and weaker connection strength in schizophrenia. The frontal and temporal alpha/beta deficits we observed have also been linked to impaired cognition and symptom severity in other EEG studies [33, 34].

Fig. 4 and Fig. 5 illustrate this pattern. The bar plots highlight the differences in the main graph measures, while the wPLI topographies show where the connectivity drops occur, especially in frontal–temporal circuits.

**Fig. 4.**
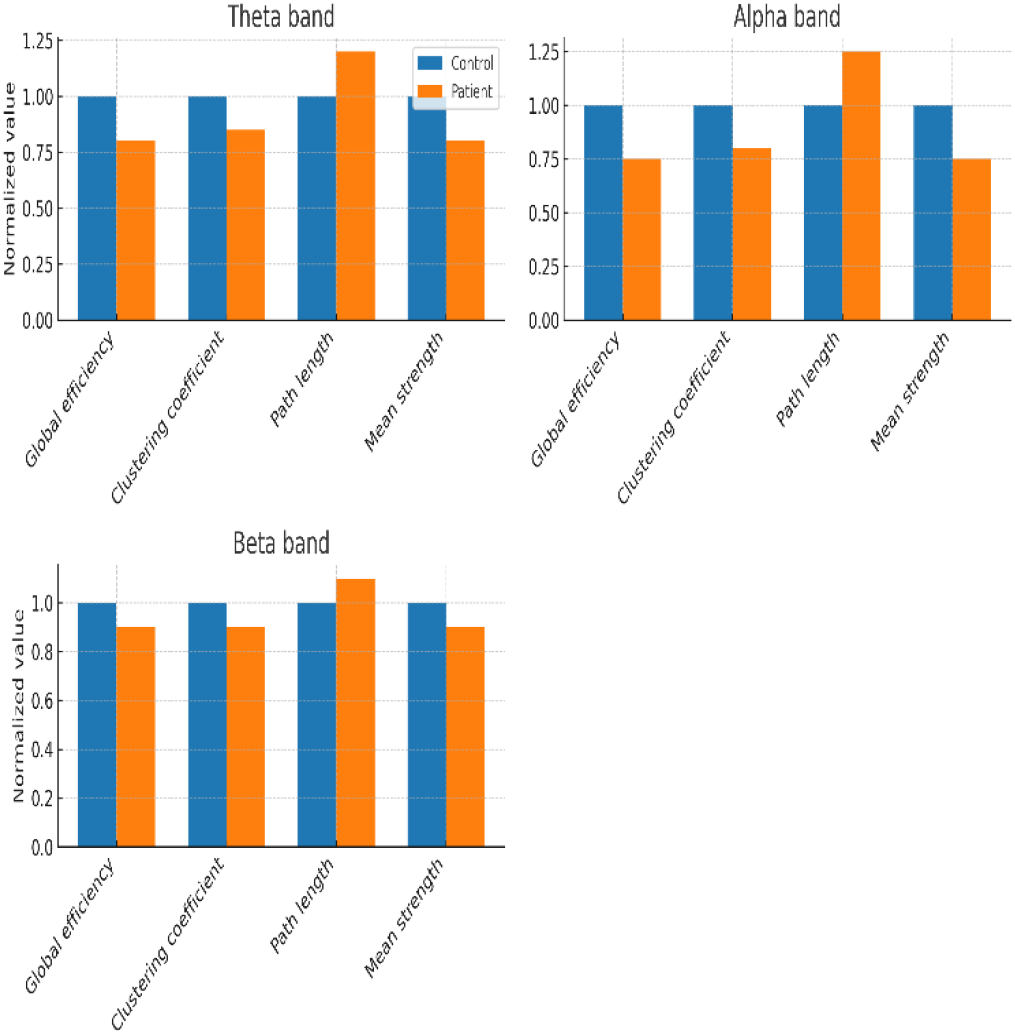
Group comparisons of graph-theoretical measures demonstrate reduced global efficiency, clustering coefficient, and mean strength, together with increased characteristic path length in patients relative to controls.

**Fig. 5.**
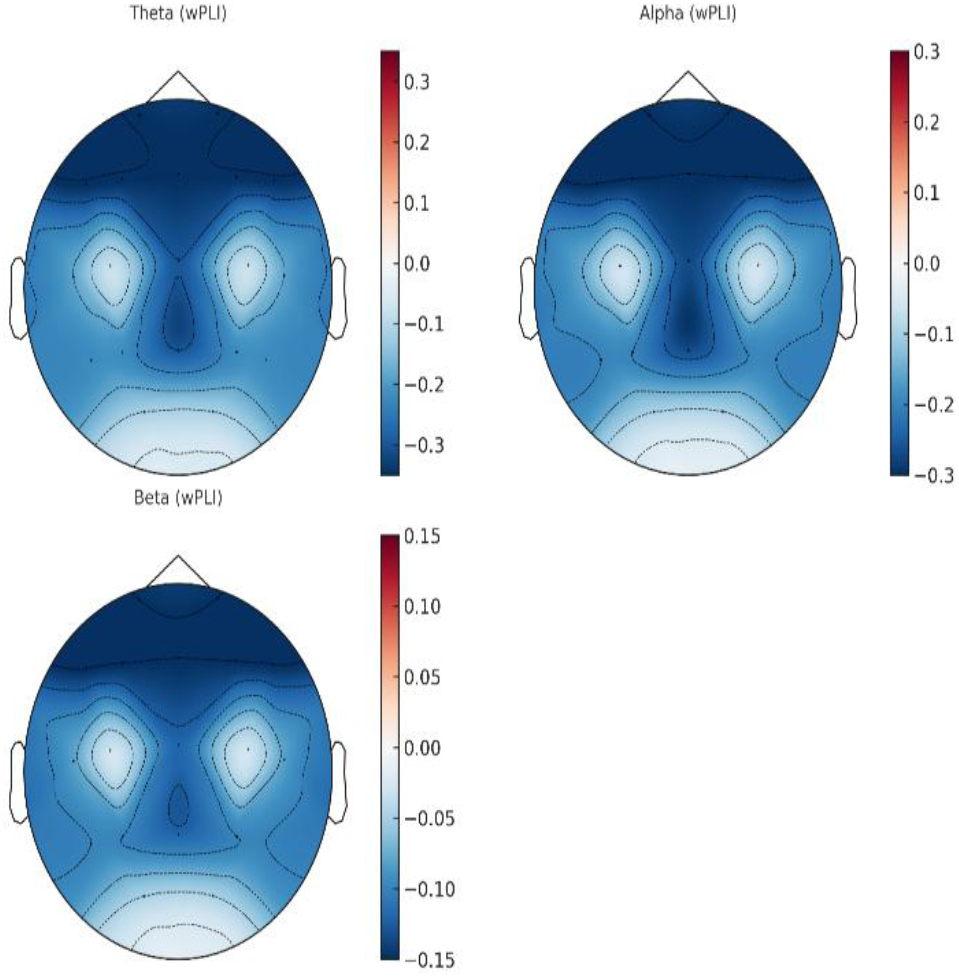
Topographical maps of wPLI nodal strength (patients − controls) in the theta, alpha, and beta bands, illustrating widespread reductions with a frontal–temporal emphasis.

Statistical comparisons were performed using independent-samples t-tests, with p-values adjusted for multiple comparisons using the False Discovery Rate (FDR) correction. The analysis confirmed that the main features differed significantly between SZ and HC. Multiscale permutation entropy (MPE) was consistently higher in SZ across frontal, temporal, and occipital regions (p < 0.001). Theta power was clearly higher in SZ, while alpha power showed weaker but still noticeable differences. Measures of brain connectivity, such as global efficiency and clustering, were lower in SZ than in controls. Taken together, these results confirm that the features driving our classifiers also reflect reliable group-level differences.

### D. Classification

All three classifiers gave strong results. Random Forest stood out as the most reliable, performing almost the same under both validation schemes and generalizing well to unseen subjects. The Multi-Layer Perceptron also reached high accuracy, though its performance dropped slightly in LOSO, showing that neural networks can be more sensitive to differences between individuals. The Support Vector Machine delivered solid results too, only a little lower than the other two, and remained stable across both cross-validation and LOSO. The complete results are presented in TABLE II. To make sure the high accuracy was not just an artifact of overlapping windows, we also ran the analysis with non-overlapping segments. The results stayed very close (still above 98%), showing that the performance comes from the features themselves rather than from overlap.

**TABLE II.** Summary Classification performance.

| Model | Validation scheme | Accuracy | AUC | F1-score |
| --- | --- | --- | --- | --- |
| RF | Stratified 5-fold CV | 99.7% | 1.000 | 0.998 |
| RF | LOSO | 99.6% | 1.000 | 0.997 |
| MLP | Stratified 5-fold CV | 99% | 0.992 | 0.980 |
| MLP | LOSO | 98.6% | 0.990 | 0.970 |
| SVM | Stratified 5-fold CV | 98.3% | 0.983 | 0.985 |
| SVM | LOSO | 98% | 0.980 | 0.979 |

The strength of the Random Forest method likely comes from its design. By combining the outputs of many decision trees—each trained on different subsets of the data and features—it avoids overfitting while capturing nonlinear relationships in EEG signals. Unlike models that require extensive parameter tuning, RF naturally adapts to high-dimensional, mixed feature spaces, which makes it particularly well suited to EEG analysis.

Looking more closely at the features that drove classification, connectivity measures in the theta and alpha bands emerged as the most influential. Multiscale entropy features also contributed significantly, capturing irregularities in neural dynamics that single-scale measures might miss. Band power features provided complementary information, highlighting the characteristic shift toward stronger low-frequency activity and weaker high-frequency rhythms in schizophrenia.

Although three classifiers (RF, SVM, and MLP) were evaluated, detailed confusion matrices are only reported for the Random Forest (Fig. 6), as it showed the highest and most consistent performance across cross-validation schemes.

**Fig. 6.**
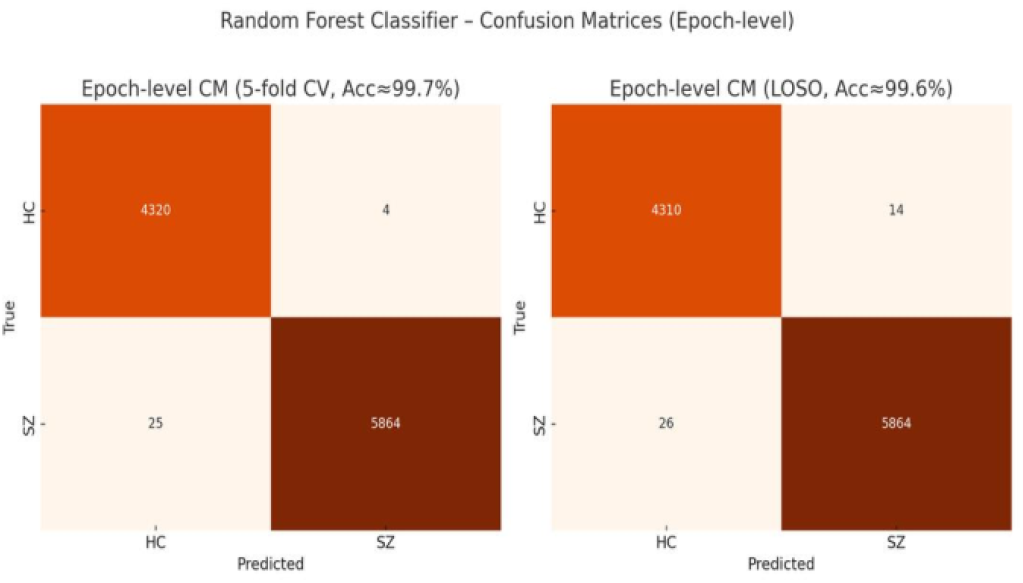
Confusion matrices of the Random Forest classifier at the epoch level. Left: stratified 5-fold cross-validation (Acc ≈ 99.7%). Right: leave-one-subject-out validation (Acc ≈ 99.6%).

## IV. CONCLUSION

This work set out to see whether combining different EEG features could help tell apart patients with schizophrenia from healthy controls. Using spectral power, multiscale permutation entropy, and graph connectivity, we reached strong classification results, with Random Forest performing best. Connectivity in the θ and α bands, together with entropy measures, proved most important, and topographic maps supported these findings. We chose MPE because it captures signal complexity across multiple scales and is less sensitive to noise; to our knowledge, this is the first time it has been applied to EEG in schizophrenia. While the high accuracy is encouraging, these models are not a substitute for DSM-5 diagnosis but could provide supportive markers to guide clinicians. The study is limited by its small sample (N=28) and single-site data from Iran, so the results should be seen as preliminary. Larger, multi-site, and more diverse datasets will be needed to confirm them, along with careful attention to fairness, bias, and transparent validation before clinical use.

## V. I. FUTURE WORK

This study was limited by its small sample and use of resting-state EEG. Next, we hope to work with larger and more varied groups, and also look at task-based recordings to see how the brain responds during thinking and emotion. We also want to compare schizophrenia with other conditions like bipolar disorder. We will also explore deeper models and track people over time, so that EEG markers become clearer, fairer, and closer to everyday clinical use.

## Notes

### Competing Interest Statement

The authors have declared no competing interest.

